# SARS-CoV-2 variants resist antibody neutralization and broaden host ACE2 usage

**DOI:** 10.1101/2021.03.09.434497

**Authors:** Ruoke Wang, Qi Zhang, Jiwan Ge, Wenlin Ren, Rui Zhang, Jun Lan, Bin Ju, Bin Su, Fengting Yu, Peng Chen, Huiyu Liao, Yingmei Feng, Xuemei Li, Xuanling Shi, Zheng Zhang, Fujie Zhang, Qiang Ding, Tong Zhang, Xinquan Wang, Linqi Zhang

## Abstract

New SARS-CoV-2 variants continue to emerge from the current global pandemic, some of which can replicate faster and with greater transmissibility and pathogenicity. In particular, UK501Y.V1 identified in UK, SA501Y.V2 in South Africa, and BR501Y.V3 in Brazil are raising serious concerns as they spread quickly and contain spike protein mutations that may facilitate escape from current antibody therapies and vaccine protection. Here, we constructed a panel of 28 SARS-CoV-2 pseudoviruses bearing single or combined mutations found in the spike protein of these three variants, as well as additional nine mutations that within or close by the major antigenic sites in the spike protein identified in the GISAID database. These pseudoviruses were tested against a panel of monoclonal antibodies (mAbs), including some approved for emergency use to treat SARS-CoV-2 infection, and convalescent patient plasma collected early in the pandemic. SA501Y.V2 pseudovirus was the most resistant, in magnitude and breadth, against mAbs and convalescent plasma, followed by BR501Y.V3, and then UK501Y.V1. This resistance hierarchy corresponds with Y144del and 242-244del mutations in the N-terminal domain as well as K417N/T, E484K and N501Y mutations in the receptor binding domain (RBD). Crystal structural analysis of RBD carrying triple K417N-E484K-N501Y mutations found in SA501Y.V2 bound with mAb P2C-1F11 revealed a molecular basis for antibody neutralization and escape. SA501Y.V2 and BR501Y.V3 also acquired substantial ability to use mouse and mink ACE2 for entry. Taken together, our results clearly demonstrate major antigenic shifts and potentially broadening the host range of SA501Y.V2 and BR501Y.V3, which pose serious challenges to our current antibody therapies and vaccine protection.

## Main text

Current neutralizing antibody and vaccine strategies against severe acute respiratory syndrome coronavirus 2 (SARS-CoV-2), the causative agent of coronavirus disease-2019 (COVID-19), were developed by targeting the prototype SARS-CoV-2 Wuhan-Hu-1 strain identified during the early phase of the pandemic (*1–12*). Several regulatory agencies recently approved a selection of these antibodies and vaccines for emergency use authorization (EUA), and their rollout to high-risk populations began late last year (*13*). For instance, Eli Lilly and Regeneron developed monoclonal neutralizing antibodies (mAbs) targeting the receptor-binding domain (RBD) of the spike (S) protein that were shown to reduce patient viral loads and COVID-19-related symptoms and hospitalizations (*14–16*). Others and we also reported the isolation and characterization several hundreds of RBD-specific mAbs from SARS-CoV-2 infected individuals (*3–5, 7, 8, 17*), and some of which are under active clinical development. Pfizer/BioNtech and Moderna developed mRNA vaccines that express a stabilized form of the S protein, which demonstrated 95% efficacy against symptomatic infection and a reduced risk of severe disease (*18, 19*). Additional vaccine modalities, such as adenovirus-based, protein subunit and inactivated vaccines, have also demonstrated reasonably good efficacy levels (*20–23*).

However, there is growing concern that the epidemic raging worldwide may be generating new SARS-CoV-2 variants that are antigenically distinct from the prototype Wuhan-Hu-1 strain, rendering current antibody and vaccine strategies ineffective (*24*). At the time of manuscript submission, approximately 455,000 SARS-CoV-2 genome sequences are listed in the GISAID database and more than 4,400 amino acid substitutions have been found in the S protein. Of the latter, 1276 are in the N-terminal domain (NTD), 596 in the RBD, 569 in the subdomain 1 and 2 (SD1-2), and 1867 in the S2 region (*25*). While these mutations are driven by the intrinsic, error-prone nature of the viral encoded RNA dependent RNA polymerase (RdRp), their survival and maintenance rely on selective advantage during natural infection, transmission, and host adaptation. For example, the S protein D614G mutation became dominant just a few months into the pandemic and was associated with greater infectivity and transmissibility, although no detectable changes in antigenicity and disease severity were identified (*26–29*). However, recently identified variants of SARS-CoV-2 UK501Y.V1 (B.1.1.7) in the UK, SA501Y.V2 (B.1.351) in South Africa, and BR501Y.V3 (P.1) in Brazil are raising serious concerns (*30–34*). They are not only rapidly displacing local SARS-CoV-2 strains, but also carry RBD mutations that are critical for interactions with the ACE2 receptor and neutralizing antibodies (*35–38*). Specifically, UK501Y.V1, SA501Y.V2 and BR501Y.V3 share the N501Y mutation previously shown to enhance binding affinity to ACE2 (*39, 40*). SA501Y.V2 and BR501Y.V3 each have three mutation sites in common within the RBD, such as K417N/T, E484K, and N501Y, which may change their antigenic profile. In addition, various deletion mutants were found in the NTD such as 69-70del and Y144del in UK501Y.V1 and 242-244del in SA501Y.V2 (*32, 36*). A few mutations in the SD1-2 region near the Furin cleavage site were also identified, such as P681H in UK501Y.V1 and A701V in SA501Y.V2. As all these mutations fall in or are proximal to major S protein antigenic sites, they may adversely affect antibody neutralization induced by natural infection or by vaccination.

To test the impact of these mutations on neutralization sensitivity, we generated a panel of 28 SARS-CoV-2 pseudoviruses corresponding with the identified mutations. These include nine mutations in the UK501Y.V1 variant, ten in the SA501Y.V2 variant, 12 in the BR501Y.V3 variant, and high prevalent single mutations in the NTD, RBD, SD1-2, and S2 domain, identified across the entire S protein in the GISAID database (Fig.1 and Fig. S1). The pseudoviruses were then subjected to neutralization tests against 12 human neutralizing mAbs, including some had received EUA and some initially isolated from SARS-CoV-2 infected patients in our laboratory, and 23 convalescent plasma collected from SARS-CoV-2 patients between January and February 2020. Our results show that SA501Y.V2 is the most resistant mAbs and convalescent plasma, followed by BR501Y.V3, and then UK501Y.V1. Such resistant hierarchy is attributed to the extent of mutations identified in NTD and RBD. Crystal structural analysis of RBD carrying triple K417N-E484K-N501Y mutations found in SA501Y.V2 bound with mAb P2C-1F11 revealed a molecular basis for antibody neutralization and escape. We also show that mutations acquired by SA501Y.V2 and BR501Y.V3 substantially improve their ability to use mouse and mink ACE2 for entry. Taken together, our results clearly demonstrate major antigenic shifts and potentially broadening the host range of SA501Y.V2 and BR501Y.V3, posing serious challenges to our current antibody therapies and vaccine protection.

**Figure 1.**
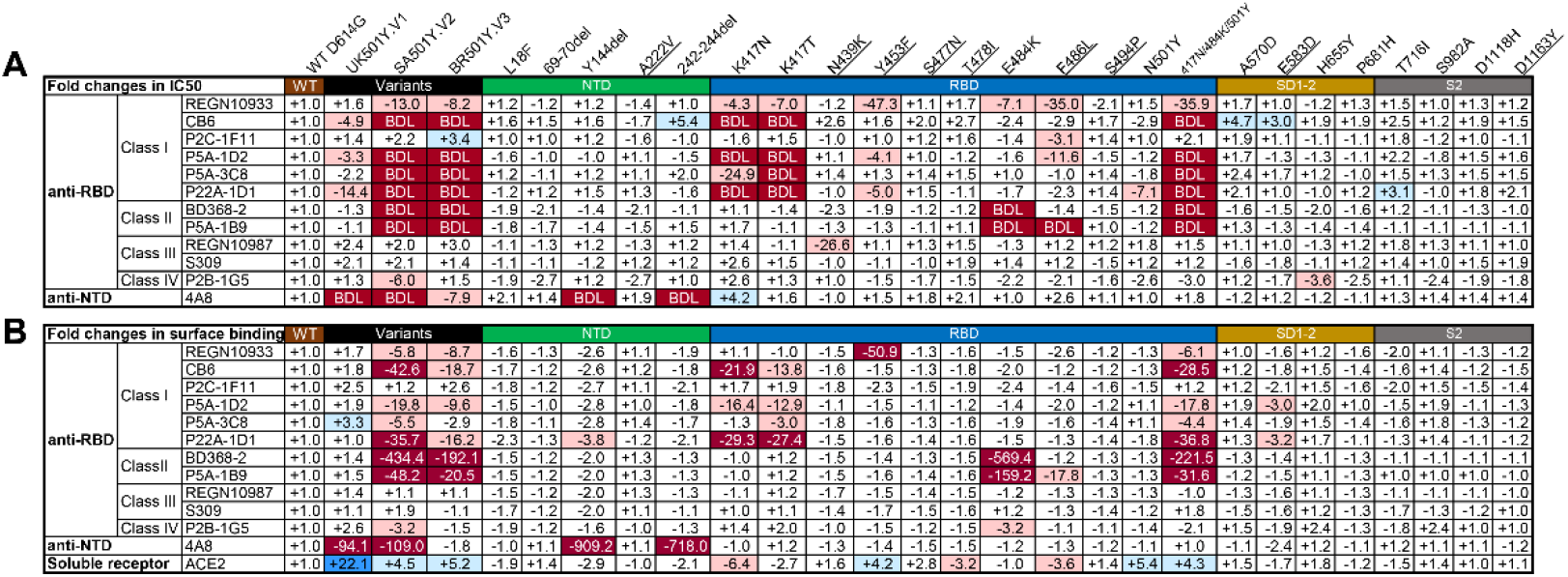
Susceptibility of SARS-CoV-2 variants to mAbs neutralization and binding. Values indicate the fold changes in (A) half-maximal inhibitory concentrations (IC_50_) and (B) mean fluorescence intensity (MFI) relative to that of WT D614G. The symbol “-” indicates increased resistance and “+” increased sensitivity. Those IC_50_ or MFI values highlighted in red, resistance increased at least threefold; in blue, sensitivity increased at least threefold; and in white, resistance or sensitivity increased less than threefold. BDL (Below Detection Limit) indicates the highest concentration of mAbs failed to reach 50% neutralization. Results were calculated from three independent experiments.

## Results

### Susceptibility of SARS-CoV-2 variant to mAb neutralization

We first evaluated the susceptibility levels of the 28 pseudoviruses to neutralization by 12 mAbs, including 11 anti-RBD and 1 anti-NTD mAbs. The anti-RBD mAbs fall into one of four major classes based on structural features affecting their mode of recognition and epitope specificity (Fig. 1 and Fig. S2) (*2, 3, 5, 17, 37, 38, 41–44*). These epitopes assignments are related to the six major sites put forward by Yuan and colleagues (*45*). The wildtype (WT) pseudovirus used throughout the analysis was the prototype Wuhan-Hu-1 strain with a D614G mutation (WT D614G). As shown in Fig. 1 and Fig. S3, SA501Y.V2 and BR501Y.V3 pseudoviruses were more resistant in terms of magnitude and breadth relative to UK501Y.V1. SA501Y.V2 and BR501Y.V3 pseudoviruses were fully resistant to almost all Class I and Class II anti-RBD mAbs except for P2C-1F11, while Class III and Class IV were largely unaffected. Many mAbs had neutralizing activity below the detection limit (BDL) even when tested at the highest concentration (1μg/mL) (Fig. 1A). The three variants were also resistant to the anti-NTD mAb 4A8, with UK501Y.V1 and SA501Y.V2 pseudoviruses showing greater resistance relative to BR501Y.V3.

Examination of the resistance patterns across the single and triple mutant pseudoviruses showed that the K417N and K417T mutations correlated with resistance to Class I mAbs, while the E484K mutation correlated with Class II (Fig. 1A). The triple mutant K417N-E484K-N501Y pseudovirus, like SA501Y.V2 and BR501Y.V3, was resistant to both Class I and II mAbs, suggesting that combination of the three RBD mutations is key in conferring complete resistance. Neutralization by the 4A8 mAb was eliminated in two NTD deletion mutant pseudoviruses, Y144del and 242-244del. These deletions were initially identified in UK501Y.V1 and SA501Y.V2 and likely contribute to their resistance profiles (*31, 32, 46*). As both deletion mutations largely fall in the NTD supersite, UK501Y.V1 and SA501Y.V2 are likely resistant to other members of the same antibody family (*8, 46–49*).

Of note, neutralization activities of mAbs approved for EUA (REGN10933, REGN10987 and CB6) or being studied for clinical use (P2C-1F11, BD368-2, S309, and P2B-1G5) were variably affected for pseudoviruses carrying mutations identified in the three variants and in the GISAID database (Fig. 1 and Fig. S3). The IC50 of REGN10933 dropped 13.0- and 8.2- fold against SA501Y.V2 and BR501Y.V3, respectively, largely due to the K417N/T and E484K mutations (Fig. 1A). Several mutations within the REGN10933 epitope, such as Y453F and F486L, were also associated with a substantial reduction in neutralization. N439K, located in the REGN10987 epitope, showed a 26.6-fold reduction in IC50. The most dramatically impacted mAb was CB6, one of the paired antibodies developed by Eli Lilly and approved for EUA, for which the neutralization against SA501Y.V2 and BR501Y.V3 pseudoviruses was below detection limit (BDL) when the highest concentration (1μg/mL) was used. The reduction and loss of neutralization are largely attributed to K417N/T (Fig. 1A). Interestingly, despite being a Class I or RBS-A antibody, P2C-1F11 was virtually unaffected by the mutations, as were the Class III and IV anti-RBD mAbs REGN10987, S309, and P2B-1G5 (Fig. 1A).

Next, we studied the binding avidity of the 12 mAbs to the 28 mutant S proteins expressed on cell surfaces. There was a strong correlation between neutralization and binding avidity (Fig. 1B and Fig. S4), indicating that compromised binding avidity is a major escape mechanism. Lastly, soluble human ACE2 appeared to improve binding to all three variants, as well as pseudoviruses bearing single N501Y and triple K417N-E484K-N501Y mutations (Fig. 1B, and Fig. S4). This finding indicates that N501Y plays a role in enhanced binding, which is consistent with earlier reports on both human and mouse ACE2 (*39, 40, 50, 51*). The single K417N mutation, however, decreased ACE2 binding by about 6.4-fold. Taken together, these results indicate that the NTD mutations Y144del and 242-244del and RBD mutations K417N/T, E484K, and N501Y confer substantial mAb resistance. The mAbs studied here with EUA would need to be optimized for the best efficacy possible against new variants.

### Structural basis for mAb neutralization and escape

We next studied the structural basis for P2C-1F11 able to maintain neutralization to the variants despite being in the Class I or RBS-A anti-RBD antibody. We determined the crystal structure of the SARS-CoV-2 RBD carrying K417N-E484K-N501Y mutations (RBD-3M) bound by P2C-1F11 at a resolution of 2.10 Å (PDB ID: 7E8M) (Table S1). Comparing with our previously reported the crystal structure of the wildtype RBD with P2C-1F11 at 3.0 Å (PDB ID:7CDI), we found the three mutations did not change the overall binding mode of P2C-1F11 to the RBD (Fig. 2A), as evidenced by 0.52 Å rmsd value for all 581 Cα atoms. However, a couple of subtle changes were identified. One is related to the interactive forces with residues 417 and the other with residue at 501. Like other Class I and RBS-A antibodies, P2C-1F11 binds to the wildtype K417 through hydrophobic and hydrogen-bond interactions. This is largely mediated by its heavy chain germline residue Y33 and Y52 (*43*). The K417N mutation would diminish these interactions but only replaced by one hydrogen bond between Y52 and mutant N417 (Fig. 2B and Table S2). Furthermore, unlike those in Class I or RBS-A, P2C-1F11 does not form salt-bridge interaction between aspartic acid (D) and K417 (Fig. 2B and 2C). The K417N mutation would therefore be less disruptive to P2C-1F11 than to those in the Class I or RBS-A such as P5A-1D2, P22A-1D1, CB6 studied here and CC12.1, CC12.3, COVA2-04 and COVA2-07 characterized elsewhere (*37, 43, 44*).

**Figure 2.**
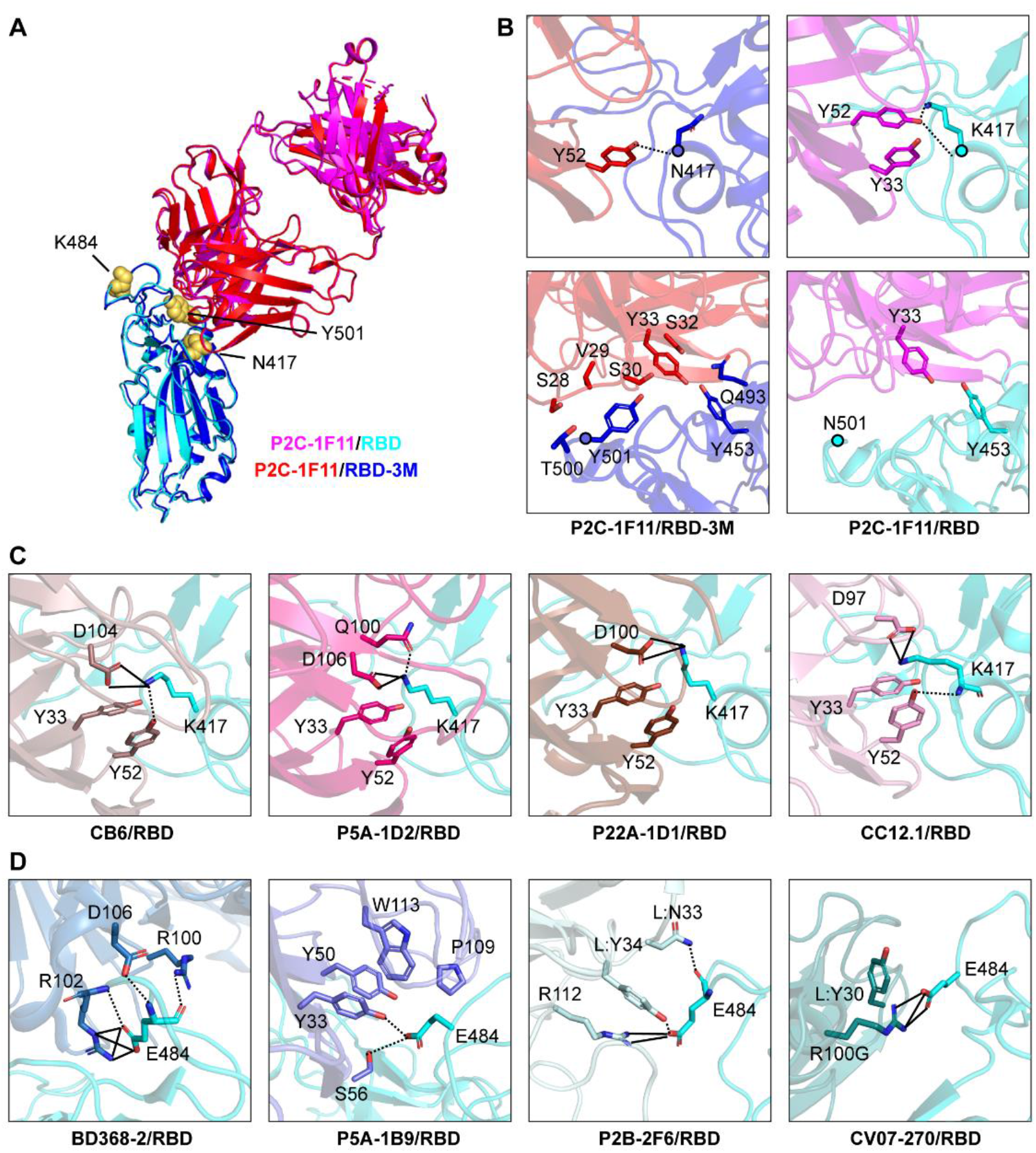
Structural basis for mAb neutralization and escape. (A) P2C-1F11/RBD-3M crystal structure superposed onto P2C-1F11/RBD crystal structure (PDB: 7CDI). P2C-1F11 is colored with magenta and RBD is colored with cyan in the P2C-1F11/RBD complex. The P2C-1F11 and RBD-3M are colored with red and blue respectively in the P2C-1F11/RBD-3M complex. The three RBD-3M mutated residues (N417, K484, and Y501) are shown as yellow-colored spheres. (B) Interactions with P2C-1F11 around RBD-3M N417 and Y501 (left panel) and wildtype RBD K417 and N501 (right panel). (C) Interactions between K417 and representative Class I IGHV3-53/3-66 antibodies CB6, P5A-1D2, P22A-1D1 and CC12.1. (D) Interactions between E484 and Class II antibodies BD368-2, P5A-1B9, P2B-2F6 and CV07-270. For panel (C) and (D), antibodies are shown with different colors; hydrogen bond and salt bridge are represented by dashed and black lines, respectively. CB6/RBD (PDB: 7C01), P5A-1D2/RBD (PDB: 7CHO), P22A-1D1/RBD (PDB: 7CHS), CC12.1/RBD (PDB: 6XC2), BD368-2/RBD (PDB: 7CHC), P5A-1B9 (PDB: 7CZX), P2B-2F6 (PDB: 7BWJ), CV07-270 (PDB:6XKP).

Structural analysis further showed that the P2C-1F11 light chain had extensive interactions with RBD-3M residues Y453, Q493, T500 and Y501 around the 501 position but had much less with the wildtype RBD residue asparagine (N) at the 501 (Fig. 2B and Table S2). The extra interactions between RBD-3M and P2C-1F11 around the Y501 may also contribute to the retained binding and neutralization of P2C-1F11 against SARS-CoV-2 variants carrying the triple K417N-E484K-N501Y mutation.

Like to the wildtype RBD, P2C-1F11 does not directly bind to E484K in the RBD-3M (Fig. 2B and Table S2). This explains E484K had no detectable impact on binding and neutralizing activity of P2C-1F11. However, the E484K mutation resulted in the complete loss of BD368-2 and P5A-1B9 neutralization (Fig. 1). Structural analysis showed that the E484 forms salt-bridge and/or hydrogen-bonding interactions with BD368-2, P5A-1B9, P2B-2F6 and CV07-270 (Fig. 2D) (*3, 41, 42, 44*). Therefore, the E484K mutation would diminish such interaction and render this class of antibodies ineffective against the SA501Y.V2 and BR501Y.V3.

### Susceptibility of SARS-CoV-2 variants to neutralization by convalescent plasma

We next studied the degree to which major variant pseudoviruses confer resistance to convalescent plasma from SARS-CoV-2 infected individuals. The plasma samples were collected from 23 patients during the early wave of the pandemic between January and February of 2020. Twelve of these patients had only mild symptoms and 11 developed severe disease. The average age was 56 with a range of 29 to 81 years old. Thirteen were men and 10 were women. For each plasma sample, eight serial dilutions were made starting from either 1:60 or 1:200 and neutralization activity was estimated based on half-maximal inhibitory dilution (ID50) and -fold changes relative to that against WT D614G pseudovirus (Fig. 3, and Fig. S5). Consistent with the findings for mAbs, SA501Y.V2 and BR501Y.V3 pseudoviruses demonstrated more resistance than UK501Y.V1 both in absolute ID50 (Fig. 3A and 3B) and -fold changes (Fig. 3B) relative to WT D614G. The average reductions in neutralization activity across the 23 plasma samples were at least 6.9-fold against SA501Y.V2, 2.4-fold against BR501Y.V3, and no significant reduction against UK501Y.V1 pseudoviruses (Fig. 3A). A complete loss in neutralization activity, indicated by below the detection limit (BDL) in Fig. 3C, was observed for 11 plasma samples against pseudovirus SA501Y.V2, four against BR501Y.V3, and none against UK501Y.V1. The remaining plasma demonstrated varying degree of reduction in neutralization potency.

**Figure 3.**
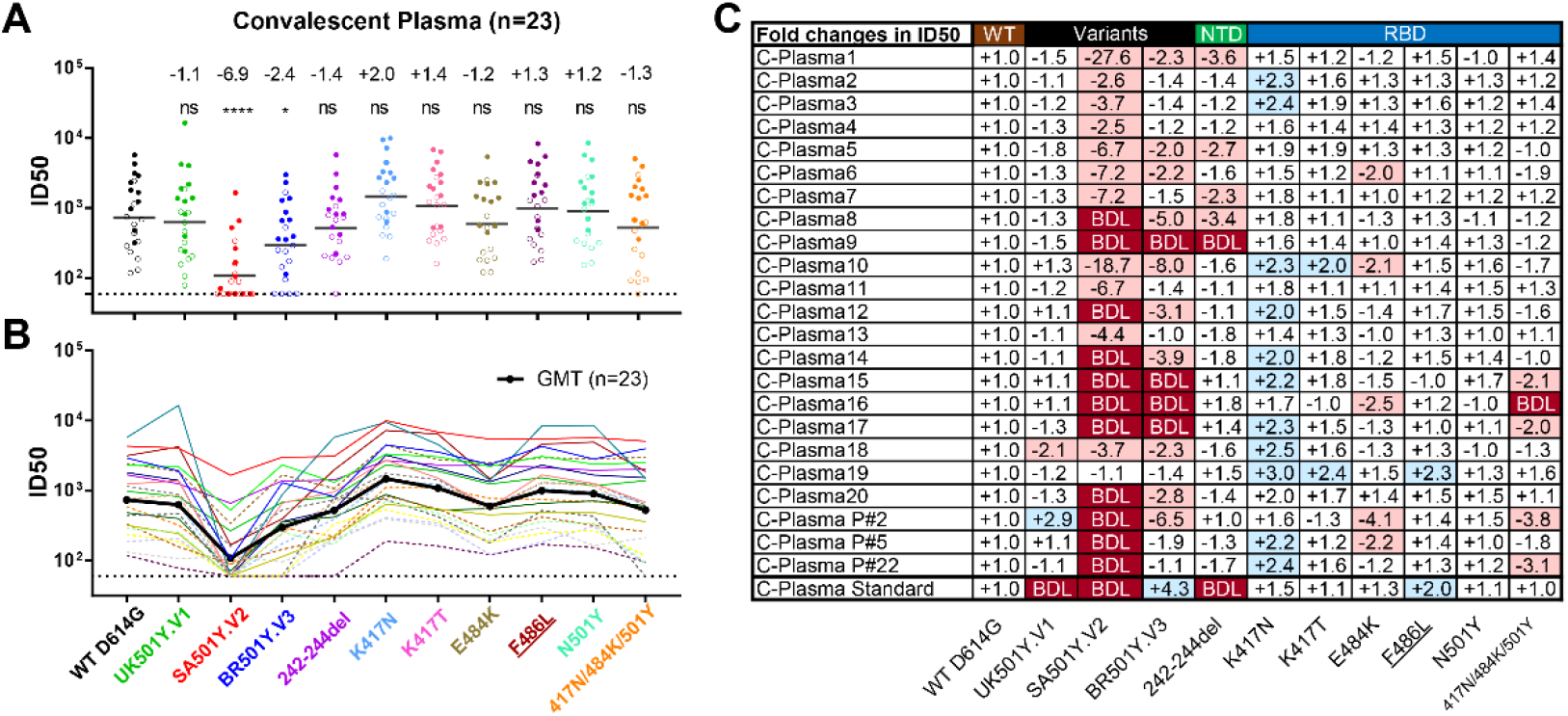
Susceptibility of SARS-CoV-2 variants to convalescent plasma neutralization. Reciprocal *plasma* dilutions (*ID50*) against SARS-CoV-2 variants were shown either by (A) colored dots or (B) colored curves, each of which represent a different convalescent plasma. The geometric mean against each variant is indicated by a back horizontal line in (A) and back curve (B). Plasma samples from mild and severe patients are indicated by empty or solid circle in (A), and dashed or solid curve in (B). The fold change in ID50 between mutant and WT D614G pseudoviruses are shown by overall average at the top in (A) or individually in (C). The symbol “-” indicates an increase in resistance while the symbol “+” indicates an increase in sensitivity. In (C), red highlights indicate a minimum twofold increase in resistance; blue a minimum twofold increase in sensitivity; and white a less than twofold change in either resistance or sensitivity. BDL (Below Detection Limit) indicates the highest concentration of plasma (1:60) failed to confer 50% neutralization. Standard plasma was obtained from the NIBSC (code: 20/136). Results were calculated from three independent experiments. *: P<0.05; and ****: P<0.0001. ns: not significant.

Examination of the resistance patterns across the single and triple mutant pseudoviruses revealed some notable patterns. In particular, the 242-244del mutation in the NTD reduced the neutralization activity more than 2-fold for C-Plasma1, C-Plasma5, C-Plasma7, C-Plasma8, and C-Plasma9, which correlates with the reduction or loss in potency observed against SA501Y.V2 pseudoviruses (Fig. 3C). The NTD supersite antibodies are likely to account for such significant reduction (*46, 48, 49*). On the other hand, the single mutant E484K and triple mutant K417N-E484K-N501Y pseudoviruses significantly reduced the neutralization activity for eight plasma samples (C-Plasma6, C-Plasma10, C-Plasma15, C-Plasma16, C-Plasma17, C-PlasmaP#2, C-PlasmaP#5, and C-PlasmaP#22), which corresponds with their diminished or loss of neutralization against SA501Y.V2 and BR501Y.V3 pseudoviruses. Antibodies targeting the receptor binding motif (RBM) likely contributed to such changes (Fig. 3C). In contrast, for pseudoviruses carrying only the K417N mutant, neutralization efficacy invariably increased for all plasma sample studied with an average 2.0-fold improvement. Similar trend was also noticed for pseudoviruses carrying only the K417T mutant (Fig. 3C). The single mutant N501Y pseudovirus, previously reported to enhance ACE2 binding (*39, 40, 52*), had limited effects on neutralizing activity of convalescent plasma.

Lastly, the plasma standard (code: 20/136) obtained from the NIBSC (United Kingdom) failed to neutralize UK501Y.V1 and SA501Y.V2 pseudoviruses, which correlates with the loss of efficacy against the 242-244del pseudovirus (Fig. 3C). This suggests that anti-NTD antibodies account for a major proportion of neutralizing activity in this sample. Collectively, these results indicate that SA501Y.V2 was the most resistant pseudovirus against the convalescent plasma tested, followed by BR501Y.V3 and then UK501Y.V1. The loss of plasma neutralizing activities is attributed, in varying degrees, to the 242-244del in the NTD, E484K, and triple K417N-E484K-N501Y mutations in the RBD, depending on in dividual plasma. Different convalescent plasma appears to respond differently to the mutated pseudoviruses, perhaps reflecting the different compositions and proportions of neutralizing antibodies in each individual generated during natural infection.

### Enhanced entry of SARS-CoV-2 variants through mouse and mink ACE2

To study the potential impact of SARS-CoV-2 variants on host range and cross-species transmission, we characterized the ability of ACE2 from nine host species to support the entry of 28 SARS-CoV-2 mutant pseudoviruses. HeLa cell lines stably expressing the ACE2 molecules were subjected to infection. The entry efficiency was measured and presented as fold-change relative to WT D614G. As shown in Fig. 4, the three variants UK501Y.V1, SA501Y.V2, and BR501Y.V3 gained substantial ability to infect HeLa mouse-ACE2, which correlated with those pseudoviruses bearing single (K417N, K417T, E484K, and N501Y) and triple (K417N-E484K-N501Y) mutations. This agrees well with the recent reports where either single N501Y or triple K417N-Q493H-N501Y mutations were found in the mouse-adapted SARS-CoV-2 strains, although the triple mutant causes more severe acute respiratory symptoms and mortality in standard laboratory mice (*50, 51*). Single N501Y mutation found in UK501Y.V1 and two of three (K417N and N501Y) found in SA501Y.V2 and BR501Y.V3 therefore likely enhanced the binding to mouse ACE2 thereby improved entry efficiency into HeLa mouse-ACE2.

**Figure 4.**
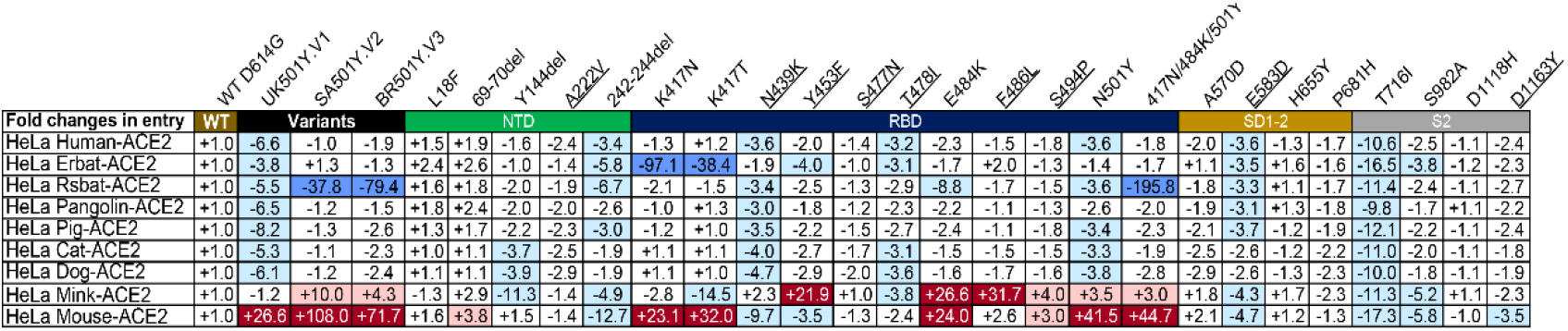
Entry efficiency of SARS-CoV-2 variants into HeLa cells expressing ACE2 from diverse host species. The values show the fold changes in luciferase activity for each indicated mutant pseudovirus variant compared to WT D614G. The symbol “+” indicates an increase in entry efficiency, while “-” indicates a decrease. Red highlights indicate at least threefold increase in efficiency; blue indicates at least threefold decrease in efficiency, while white indicates no change greater than threefold. Results were calculated from three independent experiments.

Interestingly, the single mutant E484K pseudovirus also enhanced the entry into both HeLa mouse-ACE2 and HeLa mink-ACE2. Such enhancement was correlated only with the E484K-bearing variant SA501Y.V2 and BR501Y.V3 but not with E484K-missing variant UK501Y.V1, suggesting the added and/or synergistic effect of E484K with other mutant residues in facilitating entry into these two cell lines. Furthermore, two single mutant pseudoviruses Y453F and F486L also substantially improved the entry efficiency. The two very mutations have recently been found among the mink-associated SARS-CoV-2 circulating in mink farms in Denmark, suggesting their critical role in adaptation and transmission among the mink population (*53*). Taken together, these results indicate that the three variants, particularly SA501Y.V2, and BR501Y.V3, acquired mutations in RBD that not only facilitate their escape from antibody neutralization but also potentially expand their host range to mouse and mink. Active surveillance of these variants in both human and relevant animal species would be required to minimize potential cross-species transmission.

## Discussion

In this study, we systematically evaluated the impact of mutations found in emerging SARS-CoV-2 variants UK501Y.V1, SA501Y.V2 and BR501Y.V3, as well as some single mutations within or close by the major antigenic sites in the spike protein identified in the GISAID database. We identified K417N, E484K, and N501Y mutations in the three variants that have profound consequences on antibody neutralization and interaction with mouse and mink ACE2, although mutations in NTD also impacted on antibody neutralization. It needs to be noted that our entry studies conducted on ectopically expressed mouse and mink ACE2 does not necessarily equal natural infection and transmission in the corresponding animals. However, the same K417N and/or N501Y mutations found in mouse adapted SARS-CoV-2 should raise enough concern about the potential spread of these new variants to mice and beyond. Indeed, recently identified SARS-CoV-2 variants in the mink farm in Denmark has raised another warning sign about the complexity of host range and cross-species transmission of SARS-CoV-2 variants. Rigorous and thorough monitoring of relevant animals would be required to better understand such complexity and to prevent future outbreaks.

Among the three variants, SA501Y.V2 is the most resistant against mAbs and convalescent plasma, followed by BR501Y.V3 and then UK501Y.V1. This resistance hierarchy corresponds well with genetic mutations in the NTD and RBD that led to the major antigenic changes in the spike protein. Particularly, SA501Y.V2 has both the NTD supersite mutation (242-244del) and triple K417N-E484K-N501Y RBD mutation, whereas BR501Y.V3 has only the latter and UK501Y.V1 only the former (Y144del) together with N501Y in RBD (Fig. S1). The impact of these mutations in conferring antibody resistance clearly indicates NTD and RBD are major antigenic domains on the S protein. Further study is needed to precisely estimate the relative contributions of RBD-directed and NTD-directed neutralizing antibodies to overall plasma neutralizing activity. Up till now, the most potent mAbs isolated from infected and vaccinated individuals were often dominated by those targeting the RBD and many isolated NTD mAbs failed to reach 100% potency in neutralizing activity (*3–6, 8, 46, 48, 54*). These results clearly point to greater impact of RBD antibodies in overall plasma neutralization. Nevertheless, new SARS-CoV-2 variants with an increasing number of mutations in both the NTD and RBD will challenge the efficacy of mAb therapies and vaccines.

The impacts of these mutations, in terms of mAbs, are crystal clear. SA501Y.V2 and BR501Y.V3 are resistant to neutralization by many anti-RBD and anti-NTD antibodies, including two (CB6 and REGN10933) already approved for EUA (*13, 54–56*). Most Class I mAbs studied here were disrupted by the K417N/T mutation while those in Class II by the E484K mutation. Of note, we observed that the single K417N/Y mutant tended to increase, rather than decrease, the neutralizing activity of non-Class I mAbs. As K417N/T also markedly reduced binding to soluble ACE2, this mutation may shift the balance in favor of antibody binding rather than ACE2 binding. Similar findings were also found for convalescent plasma. More importantly, P2C-1F11 currently in therapeutic development was virtually unaffected by either the single K417N/T or the triple K417N-E484K-N501Y mutation. The ability of Y52 in its heavy chain to maintain hydrogen bond with the mutant N417 while gathering more interactive forces around mutant Y501 provides the plausible structural explanations (Fig. 2 and Table S2). Considering that most Class I antibodies with heavy chain IGHV3-53/3-66 usage lost efficacy against the K417N mutation, the P2C-1F11 findings shows existence of Class I antibodies that are not affected by the mutation. This offers hope for inducing P2C-1F11-like antibodies by vaccine that can maintain strong, broadly neutralizing activity against wide range of variants, including SA501Y.V2 and BR501Y.V3.

For convalescent plasma, the most profound impact was observed on the SA501Y.V2 pseudovirus, followed BR501Y.V3 and then UK501Y.V1. The reduction or complete loss of neutralization activities were variably attributed to the 242-244del mutation in the NTD, E484K, and triple K417N-E484K-N501Y in the RBD, depending on individual profile of neutralizing antibodies. Similar reductions in the neutralization of SA501Y.V2 have also been reported elsewhere in recovered patients and individuals vaccinated with either mRNA or inactivated vaccines approved under EUA (*13, 54–56*). Although the specific levels of neutralizing antibody required to confer protection remains uncertain, reductions in antibody titers raises concerns about their protective potentials against emerging variants, particularly SA501Y.V2. Indeed, reduction in patient antibody titers have been associated with an increased likelihood of re-infection (*57*). Recent vaccine trials in South Africa showed reduced efficacy against SA501Y.V2, but maintained efficacy against wildtype strains like Wuhan-Hu-1 (*58–60*). Taken together, these results clearly demonstrate that antigenic shifts are occurring among emerging SARS-CoV-2 variants. This calls for the immediate reevaluation and update of mAb therapies and vaccines. In the long run, the goal is to develop universal therapeutic and preventive interventions that maintain efficacy against all strain mutations. Until that day comes, control over the emergence of new variants requires accelerated vaccine rollouts and ardent practice of proven public health measures.

## Acknowledgements

We acknowledge the work and contribution of all the health providers from Beijing Youan Hospital, Beijing Ditan Hospital and Shenzhen Third People’s Hospital. We also thank patients for their active participation. This study was supported by the National Key Plan for Scientific Research and Development of China (2020YFC0848800 and 2020YFC0849900), the National Natural Science Foundation Award (81530065, 91442127 and 32000661), the Science and Technology Innovation Committee of Shenzhen Municipality (202002073000002). It is also supported by funds from the COVID-19 Science and Technology Project of Beijing Hospitals Authority (YGZX-C1), Beijing Advanced Innovation Center for Structural Biology, Beijing Municipal Science and Technology Commission (Z201100005420019), Tsinghua University Scientific Research Program (20201080053 and 2020Z99CFG004), Tencent Foundation, Shuidi Foundation, TH Capital and the National Science Fund for Distinguished Young Scholars (82025022), China Postdoctoral Science Foundation (2020T130062ZX). The funders had no role in study design, data collection, data analysis, data interpretation or writing of the report. We thank the SSRF BL17U1 beamline for data collection and processing.

## Author contributions

L.Z., X.W., T.Z. conceived and designed the study. R.W., Q.Z. and J.G. performed most of the experiments with assistance from R.Z., J.L. and P.C.. J.G. and J. L. produced the RBD and ACE2 recombinant proteins, solved and analyzed the crystal structure of the antibody-RBD complex. P.C. provided an assistance in antibody production. W.R and Q.D. provided HeLa cell lines expressing ACE2 from diverse origin. T.Z., F.Z., Z.Z. and H.L. provided clinical care and management of infected patients, and particularly the recruitment and following up the study subjects. T.Z., F.Z., Z.Z, B.J., B.S., F.Y., Y.F., X.L and X.S. conducted sample collection, processing, and initial characterization. R.W., Q.Z., J.G., T. Z., X.W. and L.Z. had full access to data in the study, generated figures and tables, and take responsibility for the integrity and accuracy of the data presentation. X.W. and L.Z. wrote the manuscript. All authors reviewed and approved the final version of the manuscript.

## Competing interests

The authors have filed patent applications on some of the antibodies described.

## Data and materials availability

Data and reagents presented in the manuscript and the supplementary materials are available from the corresponding author upon reasonable request. The coordinates and structure factors files for P2C-1F11 and RBD-3M complex have been deposited to the Protein Data Bank (www.rcsb.org) with accession code 7E8M.

## Supplementary Materials

### Materials and Methods

#### Study approval

This study received approval from the Research Ethics Committee of Beijing You’an Hospital (LL-2020-039-K), Beijing Ditan Hospital (2020-019-01), and Shenzhen Third People’s Hospital (2020-084). The research was conducted in strict accordance with the rules and regulations of the Chinse government for the protection of human subjects. The study subjects agreed and signed the written informed consents for research use of their blood samples.

#### Convalescent patients and blood samples

The study enrolled a total of 23 convalescent patients aged from 29 to 81 years old, with an average of 56, infected with SARS-Cov-2 between January to February 2020. SARS-Cov-2 infection status was verified by RT-PCR of nasopharyngeal swab and chest computed tomographic scan. Of the 23 infected individuals, 13 were males and 10 were females. Twelve individuals developed only mild symptom and the remaining 11 developed severe pneumonia. Convalescent blood samples were collected during hospitalization or follow-up visits in Beijing You’an Hospital, Beijing Ditan Hospital, or Shenzhen Third People’s Hospital, within two months after symptom onset. A blood sample from a healthy control individual was also included. Blood samples were separated into plasma and peripheral blood mononuclear cells (PBMC) by Ficoll-Hypaque gradient (GE Healthcare) centrifugation. All plasma samples were heat-inactivated at 56 °C for 1h before being stored at −80 °C. PBMCs were maintained in freezing media and stored in liquid nitrogen until use.

#### Production of mAbs and Fab

P2C-1F11, P5A-1D2, P5A-3C8, P22A-1D2, P5A-1B9 and P2B-1G5 are potent neutralizing mAbs initially isolated from SARS-CoV-2 infected patients by our group and previously published (*58*). Antibodies published by other groups including REGN10933, REGN10987, CB6, S309, 4A8 and CR3022 were synthesized according to the sequences released in Protein Data Bank (PDB) (*59–63*). Antibody production was conducted by co-transfection of the heavy and light chain expression vectors into HEK 293F cells using polyethyleneimine (PEI)(Sigma). After 96h, antibodies secreted into the supernatant were captured by Protein A-Sepharose (GE Healthcare) and eluted by solution buffer Glycine pH 3.0. After further purification by gel-filtration chromatography with Superdex 200 High-Performance column (GE Healthcare), antibody concentration was determined by nanodrop 2000 Spectrophotometer (Thermo Scientific). To produce Fab fragment, purified P2C-1F11 were cleaved using Endoproteinase Lys-C (Roche) with IgG to Lys-C ratio of 4000:1 (w/w) in 10 mM EDTA, 100 mM Tris-HCl, pH 8.5 at 37°C overnight. Fc fragment were removed using Protein A-Sepharose.

#### Production of SARS-CoV-2 wild-type and variant pseudoviruses

The wildtype pseudovirus used throughout the analysis was the prototype Wuhan-Hu-1 strain (GenBank: MN908947.3) with a D614G mutation (WT D614G). The variant UK501Y.V1 (GISAID: EPI_ISL_601443) was constructed with total of 9 mutations including 69-70del, 144del, N501Y, A570D, D614G, P681H, T716I, S982A and D1118H. The variant SA501Y.V2 (GISAID: EPI_ISL_700450) was constructed with 10 mutations including L18F, D80A, D215G, 242-244del, S305T, K417N, E484K, N501Y, D614G and A701V. The variant BR501Y.V3 (GISAID: EPI_ISL_792681) was constructed with 12 mutations including L18F, T20N, P26S, D138Y, R190S, K417T, E484K, N501Y, D614G, H655Y, T1027I and V1176F. The gene of variants were synthesized In Genwiz, Inc. The single mutations identified from the GISAID database were introduced into the pcDNA3.1 vector encoding WT D614G using QuickChange site-directed mutagenesis (Agilent 210519). SARS-Cov-2 pseudoviruses were generated by co-transfecting HEK-293T cells (ATCC) with human immunodeficiency virus backbones expressing firefly luciferase (pNL4-3-R-E-luciferase) and pcDNA3.1 vector encoding either wild-type or mutated S proteins. Viral supernatant was collected 48h or 72h later, centrifuged to remove cell lysis, and stored at −80°C until use.

#### HeLa cell lines expressing ACE2 from diverse origin

HeLa cells expressing ACE2 orthologs were kindly provided by Dr. Qiang Ding at Tsinghua University School of Medicine. The cDNAs encoding ACE2 orthologs were synthesized by GenScript and cloned into pLVX-IRES-zsGreen1 vectors (Catalog No. 632187, Clontech Laboratories, Inc) with a C-terminal FLAG tag. VSV-G pseudotyped lentiviruses expressing ACE2 orthologs were produced and used to generate HeLa-ACE2 cells as previously described (*64*). For studying entry efficiency of SARS-CoV-2 variants, HeLa-ACE2 cells were added to 96 well plates, mixed with 50 ul of pseudovirus, and analyzed the luciferase activities 60 h after infection using Bright-Glo Luciferase Assay Vector System (Promega Bioscience). Absolute and fold changes between mutated and WT D614G were used to estimate the entry efficiency of SARS-CoV-2 variants.

#### mAbs and plasma neutralization using pseudoviruses

Serial dilutions of mAbs were prepared with the highest concentration of 1 μg/ml except for S309 and P2B-1G5 (10 μg/ml). Serial dilutions of convalescent plasma were prepared with the highest dilution of 1:60 except for P#2, P#5, P#22 and International Standard for anti-SARS-CoV-2 immunoglobulin (human) (NIBSC code: 20/136) where 1:200 was the initial dilution. Wild type or mutated spike pseudovirus were mixed with mAbs or plasma and incubated at 37°C for 1h. HeLa -ACE2 cells were then added into the mixture and incubated at 37°C for 60h before cell lysis for measuring luciferase-activity. Th percent of neutralization was determined by comparing with the virus control. Half-maximal inhibitory concentration (IC_50_) or dilutions (ID_50_) were calculated using GraphPad Prism 7.

#### Binding of mAb and ACE2 to cell surface-expressed wild-type and mutated S glycoprotein

The entire procedure was conducted as previously published (*65*). Specifically, HEK 293T cells were transfected with expression plasmids encoding either wild-type or mutated SARS-CoV-2 S glycoproteins, and incubated at 37 °C for 36 h. Cells were digested from the plate with trypsin and distributed onto 96-well plates. Cells were washed twice with 200 μL staining buffer (PBS with 2% heated-inactivated fetal bovine serum (FBS)) between each of the following steps. First, cells were stained with the testing mAb, S2-specific monoclonal antibody (MP Biomedicals, Singapore 08720401), or ACE2 recombinant protein at 4 °C for 30 min in 100 μL staining buffer. Then, PE-labeled anti-human IgG Fc (Biolegend 410718), anti-mouse IgG FITC (ThermoFisher Scientific A10673), or anti-his PE secondary antibody (Miltenyi 130120787) was added in 40 μL staining buffer at 4 °C for 30 min. After extensive washes, the cells were resuspended and analyzed with BD LSRFortassa (BD Biosciences, USA) and FlowJo 10 software (FlowJo, USA). The serial dilution of concentration of mAbs and ACE2 were tested and the lowest saturated concentrations were used in the assay (2 μg/ml for ACE2, 0.1 μg/ml for all mAbs except for P2B-1G5 and CR3022 where 2 μg/ml was used). HEK 293T cells with mock transfection were stained as background control. Antibody binding percentages were calculated by the ratio between mutated over wild-type MFI normalized in relative to that of S2 specific antibody. All MFI values were weighted by multiplying the number of positive cells in the selected gates.

#### Recombinant RBD and ACE2 protein

Recombinant wildtype RBD, RBD with 417N-484K-501Y mutations, and human receptor ACE2 peptidase domain were expressed using the Bac-to-Bac Baculovirus System (Invitrogen) as previously described (*66*). Specifically, SARS-CoV-2 RBD (residues Arg319 to Lys529) or ACE2 (residues Ser19 to Asp615) containing the gp67 secretion signal peptide and a C-terminal hexahistidine was inserted into pFastBac-Dual vectors (Invitrogen) and transformed into DH10 Bac component cells. The recombinant bacmid was extracted and further transfected into Sf9 cells using Cellfectin II Reagents (Invitrogen). The recombinant viruses were harvested from the transfected supernatant and amplified to generate high-titer virus stock. Viruses were then used to infect Sf9 cells for protein expression. Secreted RBD and ACE2 were harvested from the supernatant, captured by Ni-NTA Sepharose (GE Healthcare) and purified by gel filtration chromatography.

#### Crystal analysis and data collection

P2C-1F11 Fab fragments were mixed with SARS-CoV-2 RBD containing K417N-E484K-N501Y mutations at a molar ratio of 1:1.2, incubated on ice for 2 h, and further purified by gel-filtration chromatography. The purified complex was concentrated to 11 mg/mL in HBS buffer (10 mM HEPES, pH 7.2, 150 mM NaCl) for crystallization. Screening trials were performed at 18 °C. The sitting drop vapor diffusion method was used by mixing 0.2 μL of protein with 0.2 μL of reservoir solution. Crystals of RBD– Fab complexes were successfully obtained in 0.2 M Ammonium sulfate, 0.1M Tris 8.5, 12% w/v PEG 8000. Diffraction data were collected at the BL17U1 beamline of the Shanghai Synchrotron Research Facility (SSRF) and auto-processed with aquarium pipeline (*67*).

#### Structural determination and refinement

Structures were determined by the molecular replacement method using PHASER (CCP4 Program Suite) (*68*). Search models were the SARS-CoV-2 RBD structure (PDB ID: 6M0J) and the heavy and light chain variable domain structures available in the PDB with the highest sequence identities. Subsequent model building and refinement were performed using COOT and PHENIX, respectively (*69, 70*). All structural figures were generated using PyMOL and Chimera (*71, 72*).

---

**Fig. S1.**
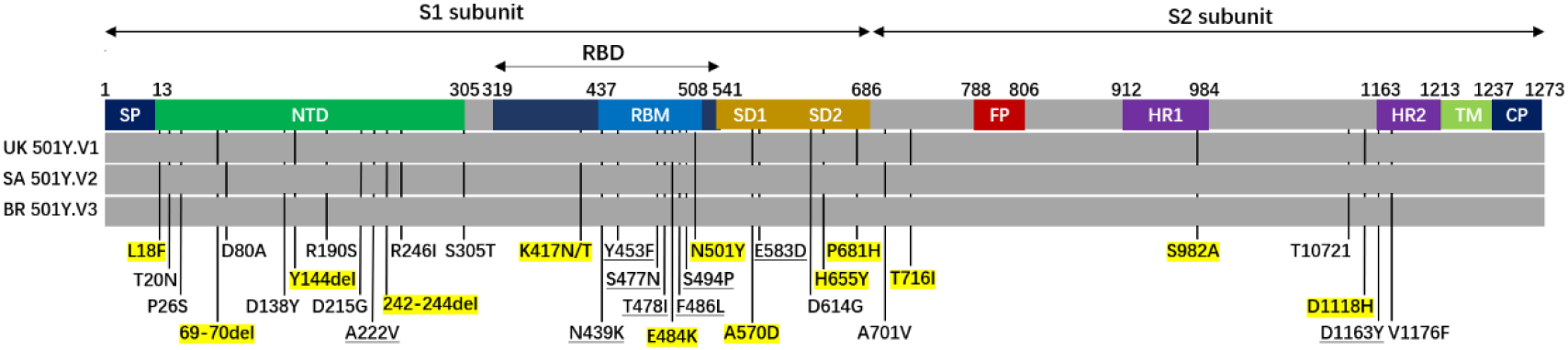
Mutant residues along the S protein identified in SARS-CoV-2 variants UK501Y.V1, SA501Y.V2, and BR501Y.V3. Each variant contains combination of mutant residues, indicated by the black vertical lines, some of which are unique while others are shared among the variants. Those highlighted in yellow represent the mutant residues studied here either singly or in combination. Nine mutant residues were underlined that were not identified in the three variants but included in the study due to their high representation in the GISAID database. SP, signal peptide; NTD, N-terminal domain; RBD, receptor binding domain; RBM, receptor binding motif; SD, subdomain; FP, fusion peptide; HR1, heptad repeat 1; HR2, heptad repeat 2; TM, transmembrane domain; and CP, cytoplasmic domain.

**Fig. S2.**
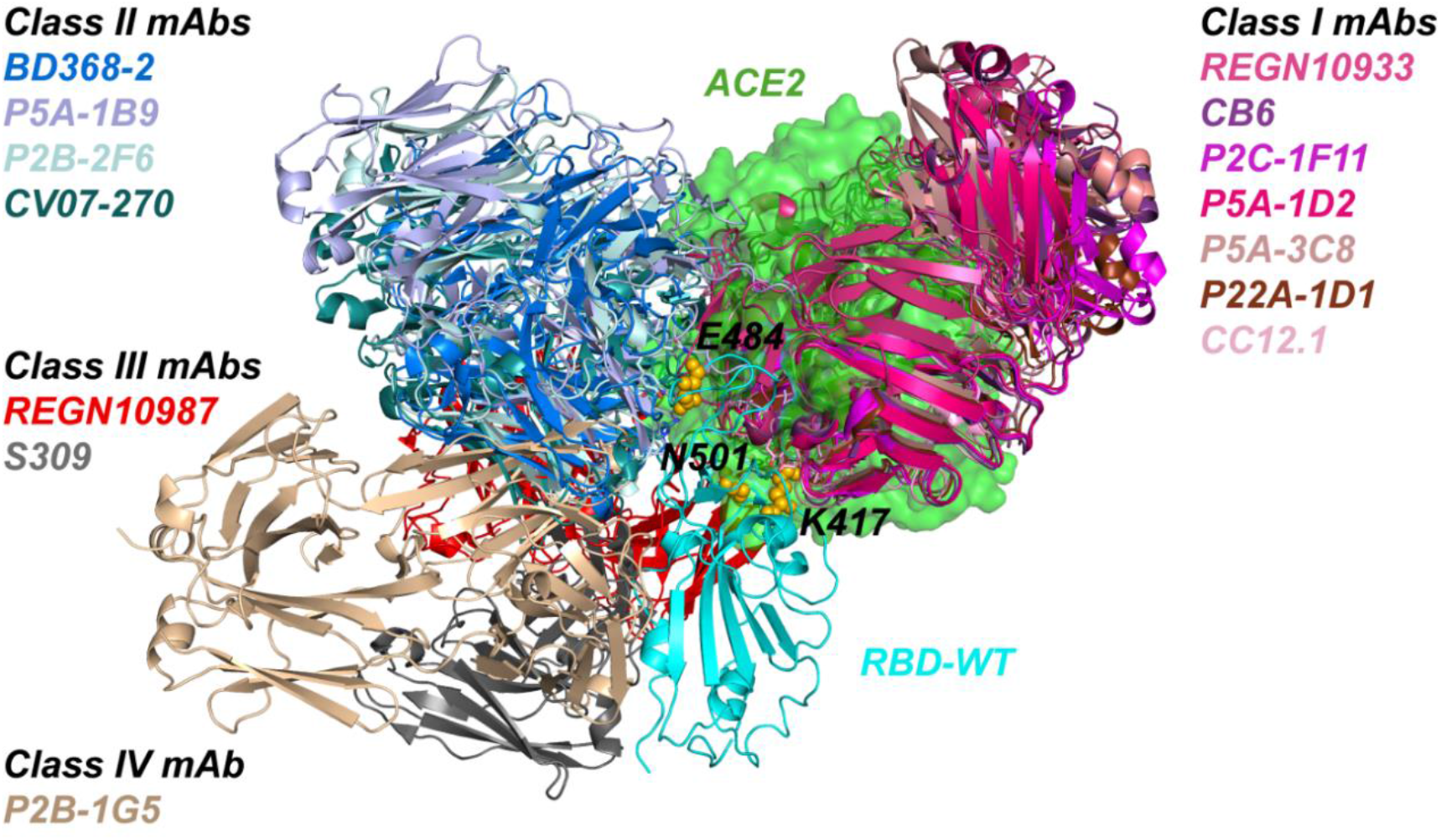
The binding mode of four classes anti-RBD antibodies. Listed antibodies are the ones studied and analyzed here. Their Fab fragments are superposed onto P2C-1F11/RBD crystal structure (PDB: 7CDI) in relative to RBD/ACE2 complex (PBD: 6M0J). P2C-1F11 is colored in magenta, RBD in cyan, and ACE2 in green. The color scheme used for the name and structure of each antibody are the same. The three residues (K417, E484, and N501) critical for binding to ACE2 and antibodies are shown as yellow-colored spheres.

**Fig. S3.**
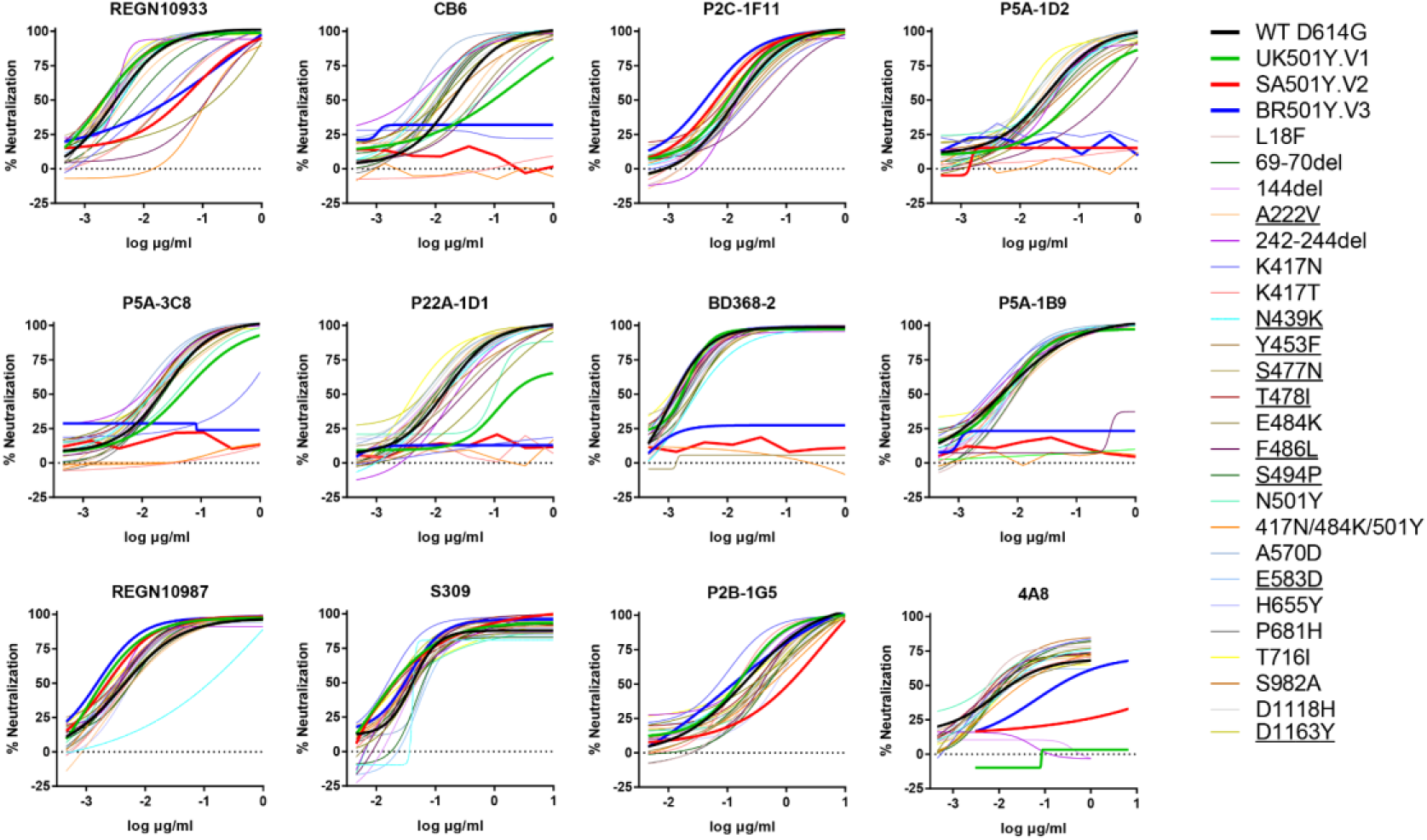
Neutralization of SARS-CoV-2 variants by each antibody. Pseudoviruses bearing the indicated mutations were tested against serial dilutions of each mAb. Neutralizing activity was defined as the percent reduction in luciferase activities compared to no antibody controls. Levels of resistance were calculated as the -fold change in IC50 between each mutant and WT D614G, as presented in Figure 1A. Results were calculated from three independent experiments.

**Fig. S4.**
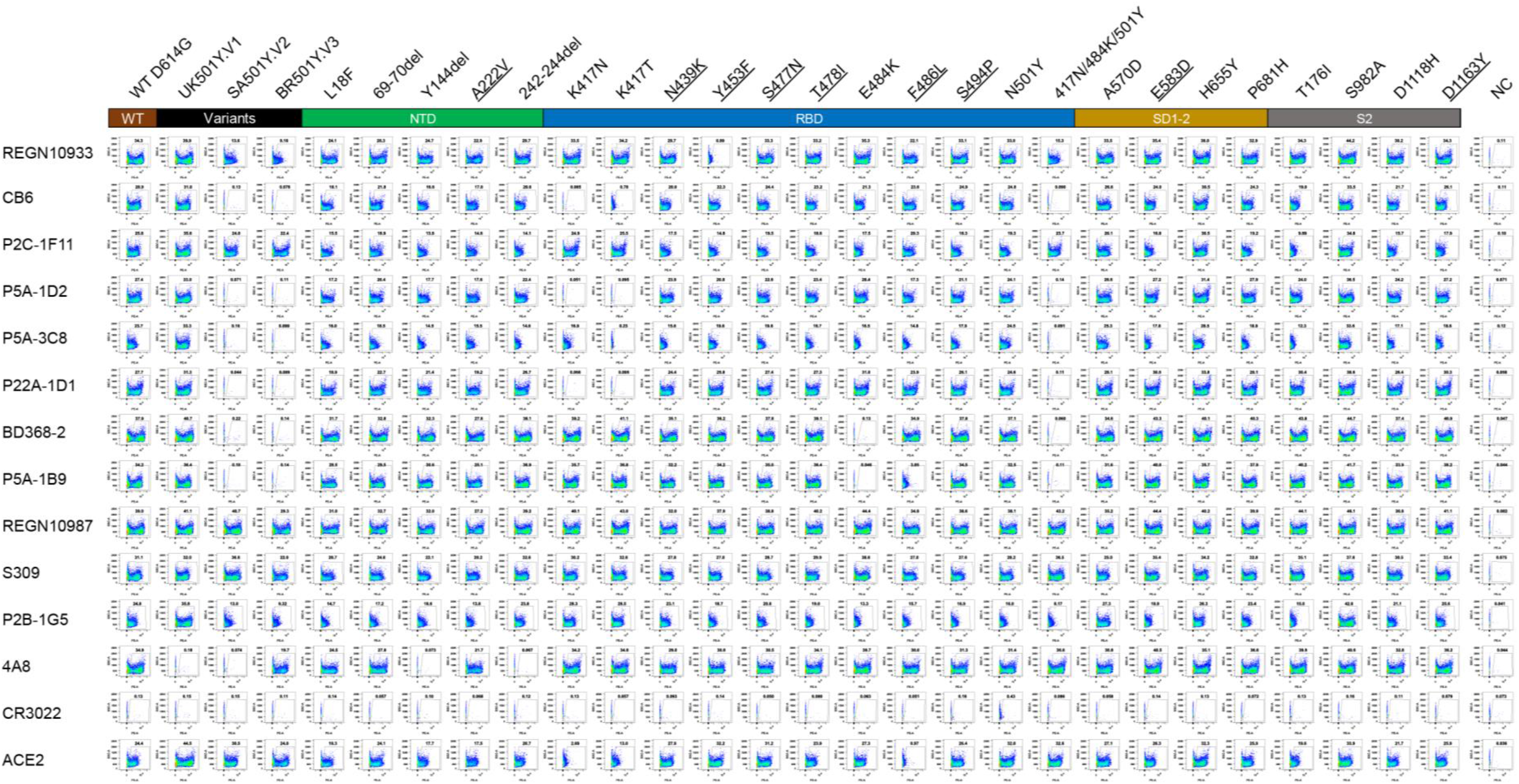
Binding to cell surface expressed SARS-CoV-2 variants by each antibody. Wildtype and mutant S proteins were expressed on the surface of HEK 293T, incubated with the mAbs or human soluble ACE2 under study, followed by staining with anti-human IgG Fc PE or anti-his PE, and analyzed by FACS. The gated cell percentages are shown. The fold changes in antibody binding, as shown in Fig. 1B, was determined by comparing the total MFI in the selected gate between S variants and WT D614G. Data shown were calculated from three independent experiments. CR3022 is a negative control antibody. NC is HEK 293T cells with mock transfection.

**Fig. S5.**
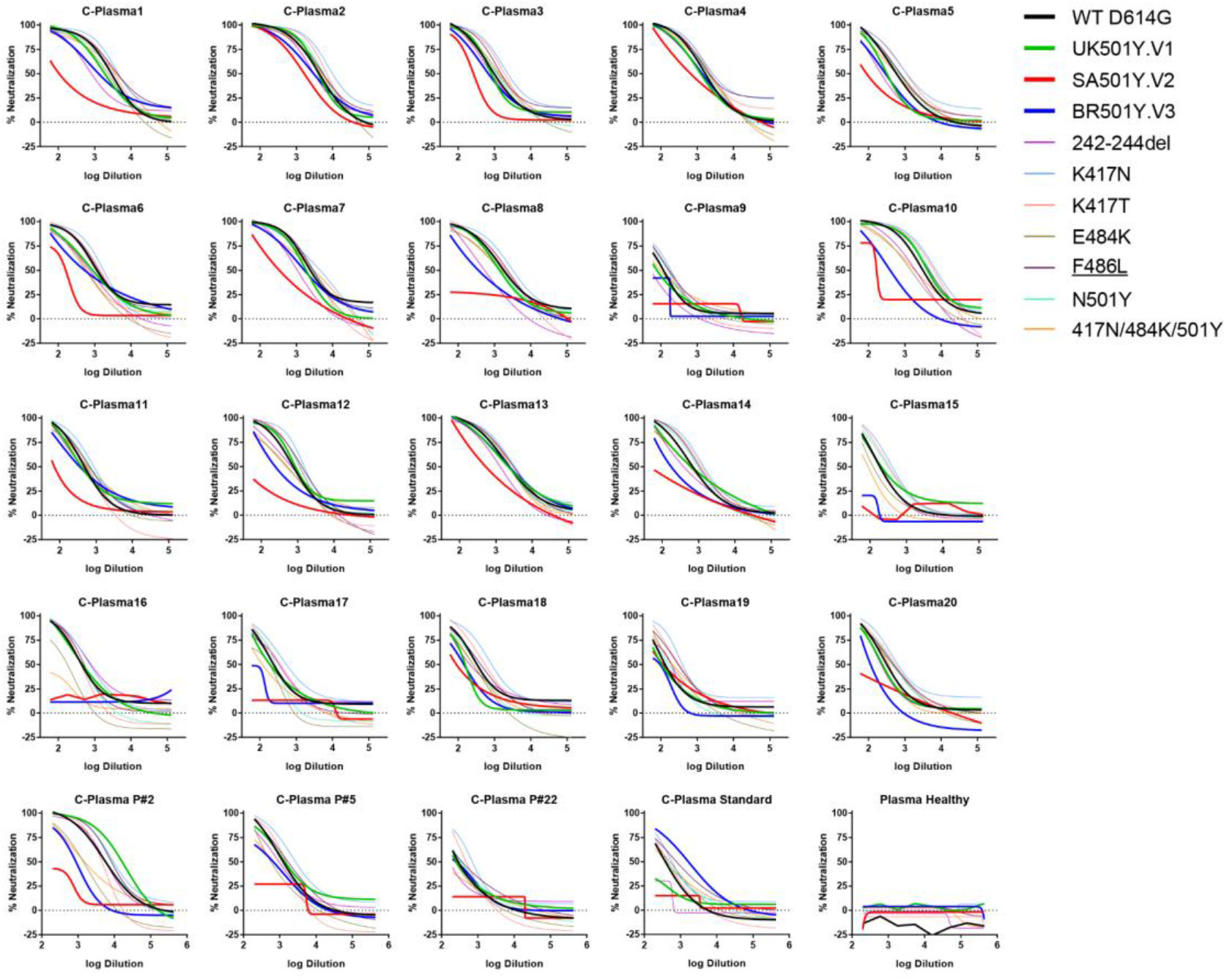
Neutralization of SARS-CoV-2 variants by each convalescent plasma. Pseudoviruses bearing the indicated mutations were tested against serial dilutions of convalescent plasma. Neutralization activity was defined as the percent reduction in luciferase activity relative to no serum control. The actual ID50 and the -fold changes between each mutant and WT D614G pseudovirus were calculated to estimate the resistance levels shown in Figure 3. Results were calculated from three independent experiments.

**Table S1.**
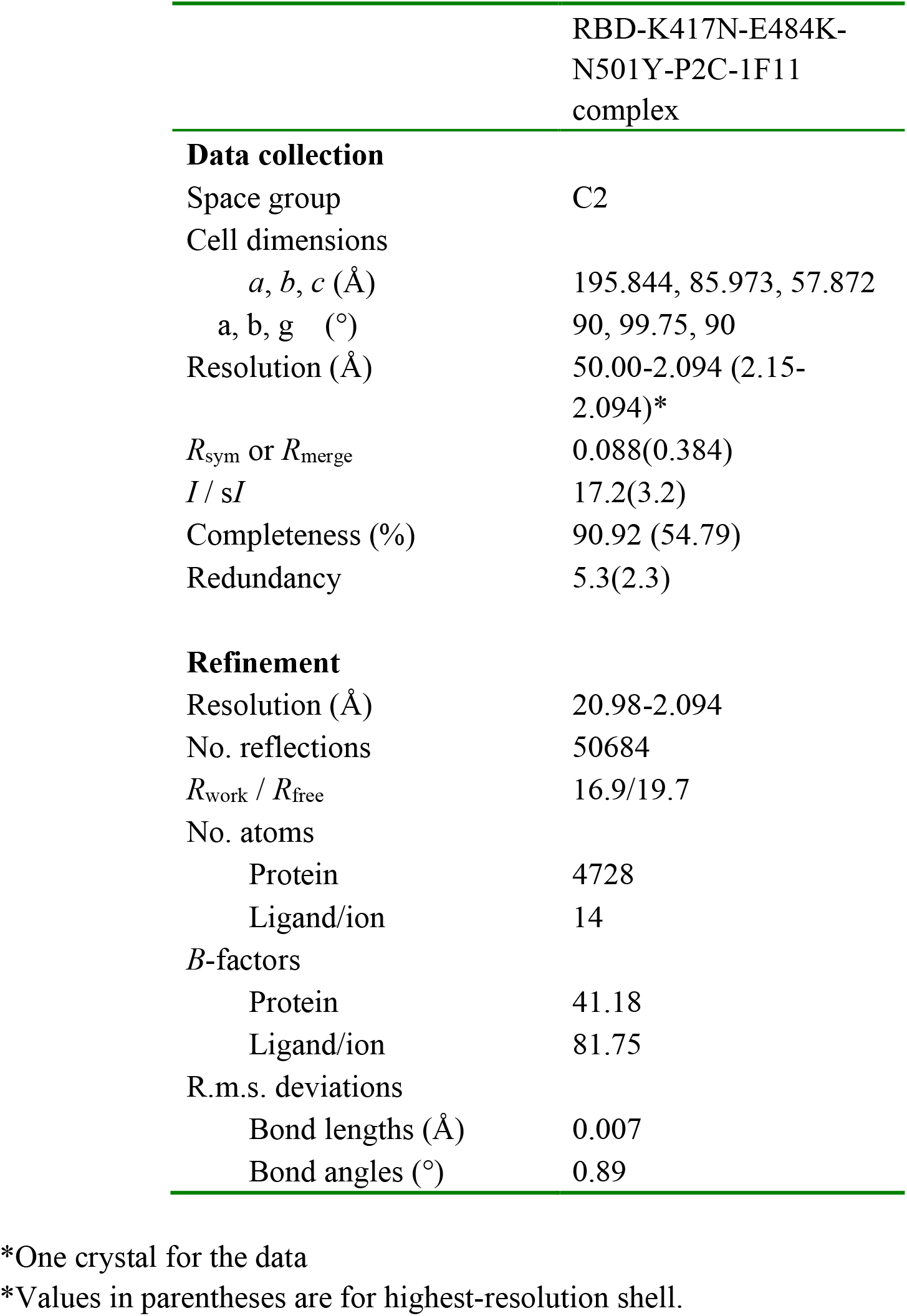
Data collection and refinement statistics (molecular replacement)

**Table S2.** Molecular interaction between P2C-1F11 and mutant RBD-K417N-E484K-N501Y

|  |  | **P2C-1F11** | **Length (Å)** | **Interactions** |
| --- | --- | --- | --- | --- |
| **P2C-1F11/RBD** | E/K417/N[N] | H/Y52/CE2[C] | 3.84 |  |
|  |  | H/Y52/OH[O] | 3.87 | Hydrogen bond |
|  | E/K417/CG[C] | H/Y52/CE2[C] | 3.85 |  |
|  |  | H/Y52/CZ[C] | 3.97 |  |
|  |  | H/Y52/OH[O] | 3.32 |  |
|  | E/K417/CE[C] | H/Y52/OH[O] | 3.69 |  |
|  | E/K417/NZ[N] | H/Y52/OH[O] | 3.19 | Hydrogen bond |
|  | E/K417/CD[C] | H/Y33/OH[O] | 4.02 |  |
|  | E/K417/CE/[C] | H/Y33/OH[O] | 4.03 |  |
| **P2C-1F11/RBD-3M** | E/N417/N[N] | H/Y52/CE2[C] | 3.82 |  |
|  |  | H/Y52/OH[O] | 3.53 | Hydrogen bond |
|  | E/Y453/CE1[C] | L/Y33/OH[O] | 3.50 |  |
|  | E/Y453/CZ[C] | L/Y33/OH[O] | 3.43 |  |
|  | E/Y453/OH[O] | L/Y33/CE1[C] | 3..58 |  |
|  |  | L/Y33/CZ[C] | 3..47 |  |
|  |  | L/Y33/OH[O] | 2.52 |  |
|  | E/Q493/CG[C] | L/Y33/OH[O] | 3.62 |  |
|  | E/Q493/NE2[N] | L/S32/CB[C] | 3.96 |  |
|  |  | L/S32/OG/[O] | 3.77 |  |
|  | E/T500/C[C] | L/S28/OG[O] | 3.90 |  |
|  | E/T500/O[O] | L/S28/CB[C] | 3.30 |  |
|  |  | L/S28/OG[O] | 3.01 |  |
|  | E/Y501/CA[C] | L/S28/OG[O] | 3.87 |  |
|  | E/Y501/CE1[C] | L/V29/O[O] | 3.79 |  |
|  | E/Y501/CZ[C] | L/S30/CA[C] | 3.94 |  |
|  | E/Y501/OH[O] | L/S30/CA[C] | 3.82 |  |
|  |  | L/S30/CB[C] | 3.85 |  |
|  |  | L/S30/OG[O] | 3.85 |  |

## Notes

### Summary of Updates

The title has been revised to better reflect the major findings of the study on both resistance to antibody neutralization and broadening of host ACE2 usage.

